# Cryogenic Soft Landing Improves Structural Preservation of Protein Complexes

**DOI:** 10.1101/2023.07.21.550105

**Authors:** Michael S. Westphall, Kenneth W. Lee, Colin Hemme, Austin Z. Salome, Keaton Mertz, Timothy Grant, Joshua J. Coon

**Author notes:** **Corresponding Author** Joshua J. Coon−Department of Chemistry, University of Wisconsin-Madison, Madison, Wisconsin 53715, United States; Department of Biomolecular Chemistry, University of Wisconsin-Madison, Madison, Wisconsin 53715, United States; Morgridge Institute for Research, Madison, Wisconsin 53715, United States;, Timothy Grant−Department of Biochemistry, University of Wisconsin-Madison, Madison, Wisconsin 53715, United States; Morgridge Institute for Research, Madison, Wisconsin 53715, United States.

## Abstract

We describe an apparatus for the cryogenic landing of particles from the ion beam of a mass spectrometer onto transmission electron microscope grids for cryo-electron microscopy. This system also allows for the controlled formation of thin films of amorphous ice on the grid surface. We demonstrate that as compared to room temperature landings, use of this cryogenic landing device greatly improves the structural preservation of deposited protein–protein complexes. Further, landing under cryogenic conditions can increase the diversity of particle orientations, allowing for improved 3D structural interpretation. Finally, we conclude that this approach allows for the direct coupling of mass spectrometry with cryo-electron microscopy.

## INTRODUCTION

Single particle cryo-electron microscopy (cryo-EM) has become a leading technology for the determination of protein structure at high resolution, even near-atomic and atomic resolution.^1^ A major bottleneck limiting the achievable resolution is the process by which protein–protein complex particles are suspended in thin films of amorphous ice—i.e., plunge freezing. With this technique—largely unchanged since its introduction forty years ago—particles often preferentially partition into the air–water interface, causing distortion upon freezing and inhibiting the random orientation that is required to solve the 3D structure.^2-4^ These issues, along with the difficulty of achieving a uniform desired ice thickness, make sample preparation a substantial obstacle and mandate considerable screening to identify particles with suitable orientations in the desired thickness of ice.^5^

Recognizing these limitations, we and others have investigated the possibility of using native mass spectrometry (nMS) for cryo-EM sample preparation as an alternative to plunge freezing. nMS, which can determine molecule weight, sub-unit stoichiometry, and collisional cross sections, is already widely utilized as a complementary structural biology tool.^6, 7^ Further, mass spectrometry (MS) requires relatively small amounts of material and has the flexibility to purify target molecules from complex mixtures. Whilst affording these orthogonal benefits of MS, deposition of particles from the ion beam of a mass spectrometer onto a graphene or amorphous carbon transmission electron microscopy (TEM) grid can in principle offer a means to bypass the above limitations of conventional cryo-EM sample preparation.

Ideally, the mass spectrometer would generate cryo-EM samples by depositing the desired number of particles onto a grid in random orientations, whilst still providing a means to coat the particles with a thin film of amorphous ice. This ice is important for the down-stream cryo-EM measurement as it provides some protection against radiation damage.^8^ There are many processes involved, however, where the particles could suffer structural degradation. First, the protein–protein complexes must be ionized, vaporized, and subjected to the numerous DC and RF fields of the mass spectrometer. Second, they must be gently deposited onto the carbon surface of the TEM grid with sufficient energy to adsorb but not so much as to cause structural damage. Third, given that deposition times typically range from one to ten minutes, particles are exposed to a high vacuum environment for long durations, where presumably any solvent that was retained will quickly be lost.

Despite these challenges, considerable efforts towards the goal of coupling MS with cryo-EM are being made by several laboratories. For example, using a modified Orbitrap system for landing cations of protein-protein complexes, we found that particles deposited onto carbon grids lacked the structural quality to enable 3D reconstructions.^9-11^ We solved this problem by using a chemical landing matrix that preserves the deposited particles so that their 3D structures could be solved to the resolution of negative stain TEM (∼ 30 A). In fact, the structures of the landed particles were virtually identical to those prepared conventionally. These matrix-landing experiments provided the best evidence to date that mass analyzed particles can retain their solution-phase structures. Unfortunately, the glycerol or polymeric matrices that we described are not directly compatible with cryo-EM. Rauschenbach et al. have constructed a device for the deposition of gas-phase cations of protein complexes onto TEM grids at room temperature followed by direct analysis with cryo-EM.^12-14^ Although the imaged particles lacked internal structure, the overall shapes were correct and assays demonstrated that the landed particles still retained biological function.^13^

We believe that amorphous ice would make the ideal landing matrix, protecting the particles from further degradation in vacuum and allowing analysis by cryo-EM. Accordingly, we have designed and constructed a modified Orbitrap mass spectrometer that contains a lander attachment capable of directing an ion beam to a TEM grid cooled to liquid nitrogen temperatures (–190 °C). Beyond offering precise temperature regulation, the instrument also permits deposition of molecular water, allowing the formation of a thin film of amorphous ice as monitored by a quartz crystal microbalance in real time. The water can be added before, during, or after particles are deposited. The system integrates a retractable ion guide so that following the cryogenic landings, the MS vacuum system can be isolated from the landing region for sample removal. Here we both describe and characterize this system for direct cryo-EM imaging of cryogenically landed cations of biomolecular complexes.

## METHODOLOGY

### Sample Purification

The purification of both β-Galactosidase and 20S proteasome core particle has recently been described by us elsewhere.^10^ The procedures are summarized below for reader convenience.

### β-Galactosidase Preparation

β-Galactosidase (from Escherichia coli, G3153-5MG) was purchased from Sigma Aldrich and prepared at 1 mg/mL in 100mM ammonium acetate. 10µL of the buffered protein was added to an additional 300µL of buffer and placed in an Amicon centrifugal filter. The sample was centrifuged (Thermo Scientific Sorvall Legend Micro 21R) at 14,000 g for 10 minutes at 4 °C. Buffer which passed through the filter was discarded and an additional 400µL of buffer was added to the sample for another round of washing using the same centrifugal settings. This wash process was repeated three times total. To obtain the final sample, the filter was inverted and centrifuged at 2000 g for 1 minute and diluted with 100mM ammonium acetate to a final concentration of roughly 1µM.

### 20S Proteasome Core Particles

Thermoplasma acidophilium expressed in E. coli BL21(DE3) was provided by Yifan Cheng (University of California, San Francisco). 100μL of the stock solution was transferred to an Amicon spin filter and centrifuged at 10,000g for 10 min at 4 °C. Three rounds of purification were done with 400 μL each of 100 mM ammonium acetate followed by centrifugation at 10,000g for 10 min at 4 °C with flow through being discarded. Final sample was obtained by inverting the Amicon filter and centrifuging at 2000g for 1 min at 4 °C.

### Mass Spectrometry and Soft Landing

Mass spectrometry experiments were performed on a modified Thermo Scientific Q-Exactive UHMR Hybrid Quadrupole-Orbitrap mass spectrometer. All instrument modifications are detailed in the Results Section. Borosilicate glass capillaries were pulled in-house using a model P-2000 laser-based micropipette puller (Sutter Instrument, CA) to an emitter inner diameter of 1 to 5 microns. A platinum wire placed within the capillary provided continuity between the mass spectrometer ESI power supply and the solution being sprayed. All full scan MS1 experiments were conducted with an ESI voltage of 1.1kV to 1.5kV, mass resolving power of 6250 at ***m/z*** 400, inlet capillary temperature of 200 °C, and no in-source trapping. For landing experiments, the Obitrap was not employed, and the ions were not stopped within the c-trap. To prevent trapping, all gas flow into the c-trap was ceased. The following DC gradient was placed on the ion optics from the inlet of the mass spectrometer to the TEM grid: Source DC offset 0V, injection flatapole 1V, inter flatapole lens 1V, bent flatapole 1V, transfer multipole 1V, C-trap entrance lens 0V, HCD field gradient 0V, HCD cell DC offset −4.5V, HCD cell exit lens 0V, and the TEM grid -1V. A wide mass filter isolation of 8,000–18,000 ***m/z*** was employed. Proteins were landed on ultra-thin carbon over lacy carbon grids (CLC400Au25-UT, EM Resolutions, UK) which were first glow discharge for 30 seconds.

### Cryo-EM Imaging and Data Processing

Soft landed grids were inserted into a Thermo Scientific Glacios microscope for imaging. Movies were collected at 200 kV using SerialEM.^15^ For β-Galactosidase, a total of 805 movies were collected for the room temperature sample and 1072 movies for the cryo-landed sample collected using a Falcon 3 camera operated in integrating mode using a pixel size of 2.58 Å. This data was collected employing a total dose of 60 e/Å^2^ over 45 frames per movie. For the 20S proteasome core particle, a total of 732 movies were collected for the room temperature sample and 1683 movies for the cryo-landed sample also using a Falcon 3 camera operated in counting mode also using a pixel size of a 2.58 **Å**. This data was collected utilizing a total dose of 89 e/Å^2^ over 67 frames per movie. All images were collected with a defocus range of 0.5 to 2.0 µm.

Collected movies were imported into ***cis***TEM 2.0 for processing.^16^ The standard workflow included motion correction and exposure filtering,^17^ followed by contrast transfer function (CTF) parameter estimation.^18^

Only micrographs with a detected CTF fit resolution better than 9 Å were kept for further processing. Next, automated particle picking, within ***cis***TEM, was employed followed by 2D classification. After selecting the best class averages, ab-inito 3D reconstruction was performed from class averages to provide starting references. Particles contained within the selected 2D class averages were carried forward for 3D auto-refinement. For both the room temperature and cryo-landed β-Galactosidase datasets, ∼50,000 particles were selected, and D2 symmetry was imposed during the ab-initio and auto-refinement steps. For the 20S proteasome, ∼14,000 particles were used in the room temperature 3D reconstruction and ∼4,000 particles were used for the cryo-landed 3D reconstruction, applying D7 symmetry in the ab-inito and auto-refinement steps. For the proteasome reconstructions, inclusion of top views caused artifacts in the final reconstructions, and angular plots demonstrated these images were largely being assigned as side views. We speculate this is due to the large droplets visible in the top view images, and in order to solve this problem only side views were carried forward into the 3D reconstruction steps. The highest resolution information included in the 3D refinement steps was 20 A for all reconstructions.

The reconstructions presented here are at relatively low resolution, particularly in the case of the room temperature samples, as such it is difficult to accurately measure the resolution. In order to provide some form of quantitative assessment of resolution, FSCs were carried out between the obtained 3D reconstructions and previously published PDBs (PDB 1F4H for β-Galactosidase and PDB 3J9I for proteosome), **Figure 1, Supporting Information**. Taking the 0.5 threshold, the measured resolutions were ∼80 A for the room temperature landed β-Galactosidase and ∼28A for the cold-landed. For the proteosome reconstructions the measured resolutions were ∼55 A for the room temperature landed sample, and ∼37A for the cold-landed.

**Figure 1.**
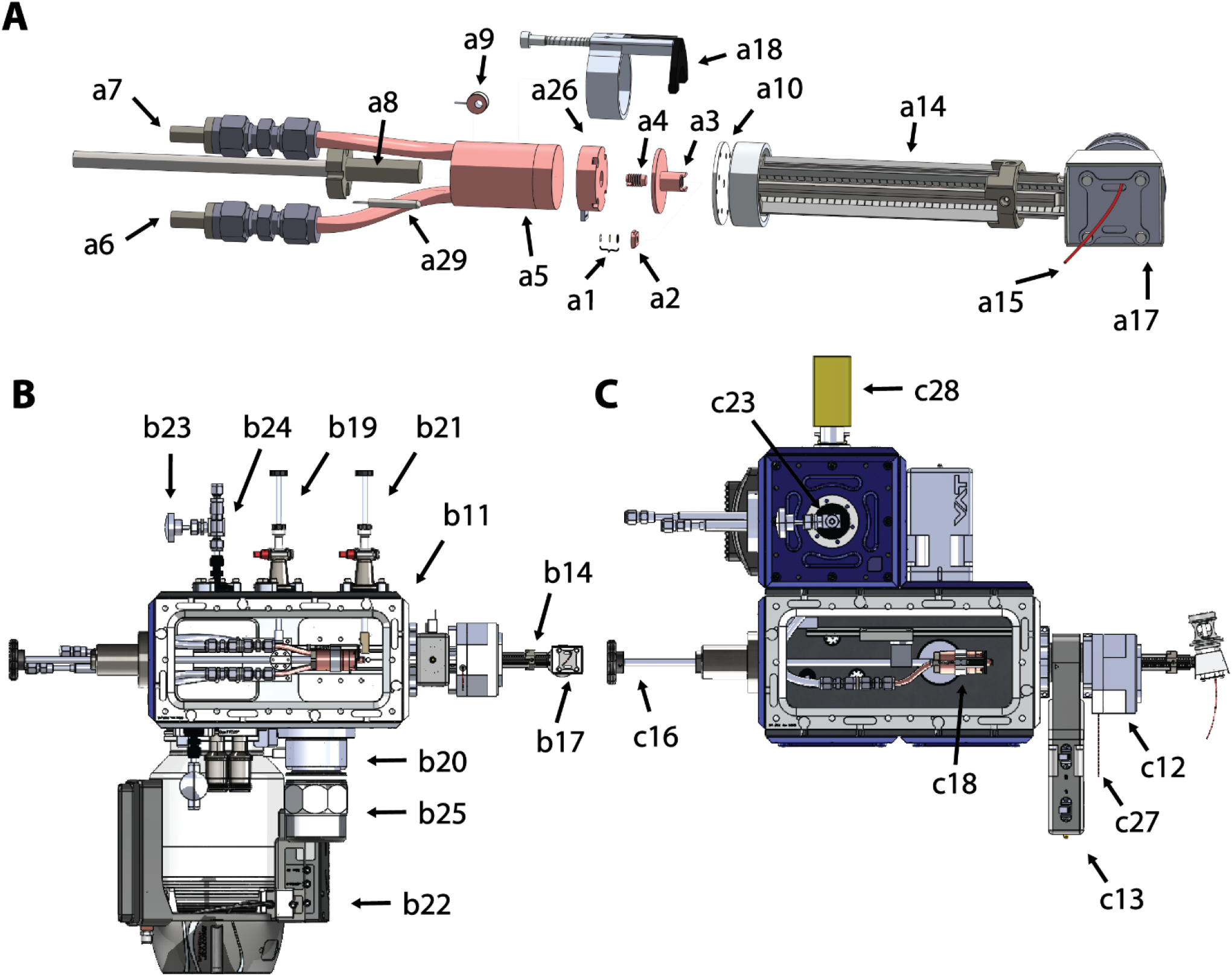
Cryogenic landing system with UHMR mass spectrometer modifications. Panel A presents an exploded view of key components developed for cryogenic soft landings. Panel B presents a side view of the cryo-lander system with the side cover from the vacuum chamber removed for clarity. Panel C presents a top view of the landing assembly with the top vacuum cover removed for clarity. Components are labeled with numbers **1** through **28** and a prefix of **a** (exploded), **b** (side), or **c** (top) to indicate the corresponding view. Individual components, depending on their position, may be detectable in multiple views; e.g., component 14 is represented in a14 and b14. The components are as follows: 1) autogrid assembly, 2) autogrid top plate, 3) autogrid transfer assembly body, 4) copper set screw, 5) copper heatsink, 6) PEEK liquid nitrogen feed, 7) PEEK liquid nitrogen and nitrogen vapor return, 8) PEEK cold block insulator, 9) cryo-temperature sensor, 10) HCD cell exit lens, 11) cryo-lander vacuum chamber, 12) modified vacuum housing of UHMR, 13) vacuum gate, 14) exposed HCD cell, 15) nitrogen gas feed to UHMR C-trap, 16) linear actuator for insertion of TEM grid holder into mass spectrometer, 17) UHMR C-trap, 18) spring loaded clamp, 19) linear actuator for TEM grid holder release, 20) vacuum gate, 21) linear actuator for TEM grid cover, 22) turbo molecular pump, 23) vent gas inlet valve, 24) pressure relief valve, 25) Dewar, 26) micro balance, 27) PEEK water vapor inlet line, and 28) vacuum gauge.

## RESULTS

To test the hypothesis that cations of protein–protein complexes could be preserved and prepared for cryo-EM using MS, we constructed the landing apparatus shown in **Figure 1**. The foremost design consideration for this device was to ensure sufficient thermal contact to the TEM grid so that even under the vacuum environment, heat could be removed fast enough to allow formation and maintenance of amorphous ice—at least 10^6^ degrees Kelvin per second.^19^ Since the ultimate destination of the TEM grids was a Thermo Fisher cryo-EM system outfitted with an autoloader, each grid was initially mounted in an autogrid ring and c-clip (Thermo Fisher Scientific part numbers 1036173 and 1036171, respectively). To interface with the autogrid assembly, we designed a holder which not only enabled excellent thermal contact to the grid but also provided sufficient thermal mass to prevent rapid temperature swings at the grid. **Figure 1** presents an exploded view of the grid transfer assembly in **a1 through a4**. Note the total mass of a single TEM grid is ∼1 mg, whilst the grid transfer assembly weighs in at 8 grams—a ∼10^4^-fold difference. Aside from the mass difference, the grid transfer assembly affords quick energy transfer to the grid by sandwiching the autogrid assembly (**Figure 1, a1**) with a threaded copper set screw (**Figure 1, a4**) against the holder top plate (**Figure 1, a2**) that slides through a slot in the grid transfer assembly body (**Figure 1, a3**).

Having developed a grid holding device, we turned our attention to construction of a cooling apparatus that would precisely regulate the temperature of the grid transfer assembly to as low as –190 °C at pressures ranging from atmosphere to vacuum. The central framework of the temperature regulator is a copper heat sink that has within it an open channel for the passage of nitrogen (**Figure 1, a5**). To isolate the heat sink thermally and electrically, we fabricated PEEK bushings to go between the cold block and linear actuating rod (**Figure 1, a8**) as well as at the attachment points of the liquid nitrogen feed and return lines (**Figure 1, a6 and a7**). Also attached to the temperature regulator’s copper heat sink are a resistive heating element (HT15W, ThorLabs, Newton, NJ) (**Figure 1, a29)** to adjust temperature and a LakeShore DT-470 series silicon diode (Lake Shore Cryotronics, Westerville, OH) (**Figure 1, a9**) to track temperature. Finally, the temperature regulator module is electrically connected to an RBD 9103 autoranging picoammeter (RBD instruments, Bend, OR) so that ions landing on the TEM grid surface can be detected.

An additional objective is to embed particles within amorphous ice. An unintended advantage of having a small vacuum leak to atmosphere was the introduction of water vapor which condensed and froze on the grid during particle landing. As shown below, this provided additional protection to the particles during landing and imaging. This arrangement is not optimal as there is no means to control the rate of ice buildup or prevent other contaminants from entering the system. To obtain a more controlled deposition, we added a third module which offers the ability to calibrate the rate of ice formation. This component is inserted between the copper heat sink and the grid transfer assembly and is based on a cryogenically cooled quartz crystal microbalance constructed in house (**Figure 1, a26**). Fashioned with a copper housing, the microbalance forms an extension of the cold block so that the grid transfer assembly continues to be sufficiently cooled. Briefly, as vapor or ice condenses onto the cryogenic temperature quartz surface, the frequency at which it oscillates changes. The change in frequency directly corresponds to the thickness of ice that has built up on the surface of the microbalance. These frequency changes are sufficiently sensitive to permit preparation of ice thickness to within a single nanometer. Note that the quartz crystal microbalance device is used to calibrate the rate of water vapor introduction prior to ion beam deposition and while the grid transfer assembly is removed. Once this calibration is complete, the grid transfer assembly may be put in place for landing experiments. Water vapor for ice formation is introduced from the head space above water (American Society for Testing and Materials Type 1) contained within a glass vial which was previously evacuated by the mass spectrometer pumping system. The vapor is introduced into the HCD cell vacuum chamber through PEEK tubing (**Figure 1, c27**) and regulated with a pin valve.

The cryo-lander depicted in Figure 1B is housed in its own vacuum chamber (**Figure 1, b11**) and aligns the TEM grid with the last ion optic of the mass spectrometer (i.e., the exit lens of the HCD cell) (**Figure 1, a10**). We have modified a Quadrupole Orbitrap mass spectrometer (Thermo Fisher Scientific Orbitrap UHMR) to accept this device by shortening the vacuum chamber which houses the HCD cell (**Figure 1, c12**) to minimize the distance between the exit lens of the HCD cell and the vacuum gate that separates the cryo-lander from the mass spectrometer. To reduce background contaminants that could form on the cryogenically cooled grid surface, we lowered the overall operating pressure of the central and rear regions of the mass spectrometer. To this end, we modified the original encased HCD cell by removing the outer enclosure and ceased the flow of nitrogen into the cell (**Figure 1, a14**). Doing so reduced pressure from 10^−4^ bar to 10^−6^ bar, in effect removing the function of mass analysis in the Orbitrap, as C-Trap and HCD cell pressures are too low to stop ions. To restore the ability to put ions in the Orbitrap, we ran a new gas line into the C-Trap (**Figure 1, a15**), pressurizing the C-Trap (**Figure 1, a&b17**) with nitrogen only when mass analysis is required, but stopping the gas flow for all landing experiments. Finally, the HCD cell RF and DC, axial DC gradient, and accompanying exit lens were operated using Modular Intelligent Power Sources (GAA Custom Electronics, Kennewick, WA).

This setup enables the process of cryo-landing. To load the grid, we vent the cryo-lander and open the device by first retracting the horizontal actuator (**Figure 1, c16**) until the plunger of the spring-loaded clamp (**Figure 1, a&c18**) intersects with the fully inserted centermost vertical linear actuator (**Figure 1, b19**). We then open the sample transfer vacuum gate (**Figure 1, b20**) to insert the grid transfer assembly through the bottom of the vacuum chamber up to a spot between the cold block and spring-loaded clamp, move the horizontal actuator forward 1cm to close the gripper, and hold the grid transfer assembly firmly in place. With the grid transfer assembly in place, we retract the centermost vertical actuator and insert the outermost vertical linear actuator (**Figure 1, b21**), which places a PEEK cover over the grid to prevent contamination during the cooling process. To return to vacuum we utilize a turbo molecular pump (Turbovac 350i, Leybold, Cologne, DE) (**Figure 1, b22**) to achieve a vacuum of 1×10^−5^ Torr. Once at vacuum the flow of liquid nitrogen through the cold block begins. Control of the nitrogen flow and overall temperature regulation is provided though a cryogenic temperature controller (CTC100, Stanford Research Systems, Sunnyvale, CA). After the cryo-lander module reaches –190 °C, we raise the outermost vertical linear actuator, releasing the grid transfer assembly cover, open the vacuum gate between the lander and mass spectrometer (**Figure 1, c13**), and using the horizontal actuator insert the module (**Figure 1, C16**) into the mass spectrometer such that the TEM grid is within one to two millimeters of the exit lens of the HCD cell. Note, the mass spectrometer ion beam has previously been tuned and directed in the Orbitrap and cryo-lander systems. Then, the ion current impinging on the grid is checked using the picoammeter to ensure the ion current is around 0.1 nanoamps. Landing typically takes 10 to 20 minutes, with the grid in place behind the HCD cell.

Once landing is complete, the extraction process is as follows. The horizontal linear actuator is retracted and the outermost vertical actuator is lowered to return the cover to the front of the grid. Next, the vacuum gate between the lander and mass spectrometer is closed, separating the lander from the mass spectrometer, and the cryo-lander pumping system is turned off. At the same time, the nitrogen gas inlet valve (**Figure 1, b&c23**) is opened to flood the cryo-lander chamber with nitrogen vapor from the headspace of a liquid nitrogen Dewar. Once the pressure inside this chamber reaches 1 psi, a safety valve (**Figure 1, b24**) opens to prevent further buildup of pressure. From this point on, the nitrogen vapor is flowing, and the liquid nitrogen continues to flow through the cold block to provide cooling. Continuous nitrogen vapor flow keeps the chamber at a positive pressure, preventing contamination from atmospheric water vapor. We now place a small Dewar (**Figure 1, b25**) of liquid nitrogen below the sample transfer vacuum gate, which provides a continuous vapor of nitrogen pushing out atmosphere in the region and a pool of liquid nitrogen in which the grid transfer assembly will be deposited. We next lower the centermost vertical linear actuator; open the sample transfer vacuum gate; and then retract the horizontal actuator until the plunger of the spring-loaded clamp intersects with the center vertical actuator, opening the clamp and releasing the grid transfer assembly, which drops into the liquid nitrogen with the sample preserved at cryogenic temperature.

To test the cryo-lander, we chose to examine β-Galactosidase, as its structure is well understood and it serves as a common cryo-EM reference complex. We performed native electrospray of a β-Galactosidase solution in ammonium acetate buffer. Using the setup described above, except set at room temperature, we deposited cations (from the entire charge envelope, z ∼42–48) of β-Galactosidase onto amorphous carbon-coated TEM grids for 15 minutes with an average energy of 3–4 eV per charge. The grid was removed as described above and kept submersed in liquid nitrogen until it was placed into a 200 keV Thermo Fisher Scientific Glacios Cryo-EM system.

Figure 2. depicts a comparison between the room temperature landing and the cryogenic landing using the device described above. Panels A and B present representative micrographs of the room temperature and cryo-landed particles, respectively. It is immediately evident that the particles shown in Panel A are free of ice, and thus the contrast of the particles is high. In Panel B, by contrast, there appears to be a thin film of what we suppose to be ice in addition to the particles, which therefore exhibit lower contrast. The formation of this thin film was likely due to contaminant water vapor in the vacuum system, which condensed whilst the particles were deposited. Note we had not yet utilized the ice formation module of the instrument when these data were collected. Results using that method of ice making are detailed below. Panels C and D show 2D class averages of particles from the respective landing conditions. The class averages from the room temperature landing lack features, appear to be damaged, and appear to have a strong preferential orientation. By contrast, the 2D class averages obtained from the cryo-landing demonstrate more features and a much wider range of orientations (**Figure 3**). Similarly, the 3D reconstruction in Panel F shows better conserved features than the reconstruction derived from room temperature landing in Panel E. Indeed, the Panel E reconstruction is so featureless as to at best represent only the molecular envelop of β-Galactosidase. In the 3D reconstruction derived from cryo-landing, however, the contour generally follows the known structure of β-Galactosidase (Panel G).

Next, we sought to test our ability to obtain a more controlled water deposition and ice formation as well as to test the effect of cryo-landing on a larger complex, 20S proteosome core particles. We first coated the cryogenically cooled TEM grid with ∼10 nm of amorphous ice by opening the water vapor valve (see above) and waiting ∼5 minutes (i.e., the device was calibrated using the quartz crystal microbalance for a deposition rate of ∼2 nm ice per minute). The water vapor valve was then closed and immediately followed by deposition of the entire charge envelope of proteosome cations (base peak z = 63) for 15 minutes. At this point the TEM grid was removed from the instrument as described above and placed into the electron microscope and kept cold the entire time. Note we also collected a room temperature control deposition of the same proteasome cations.

**Figure 2.**
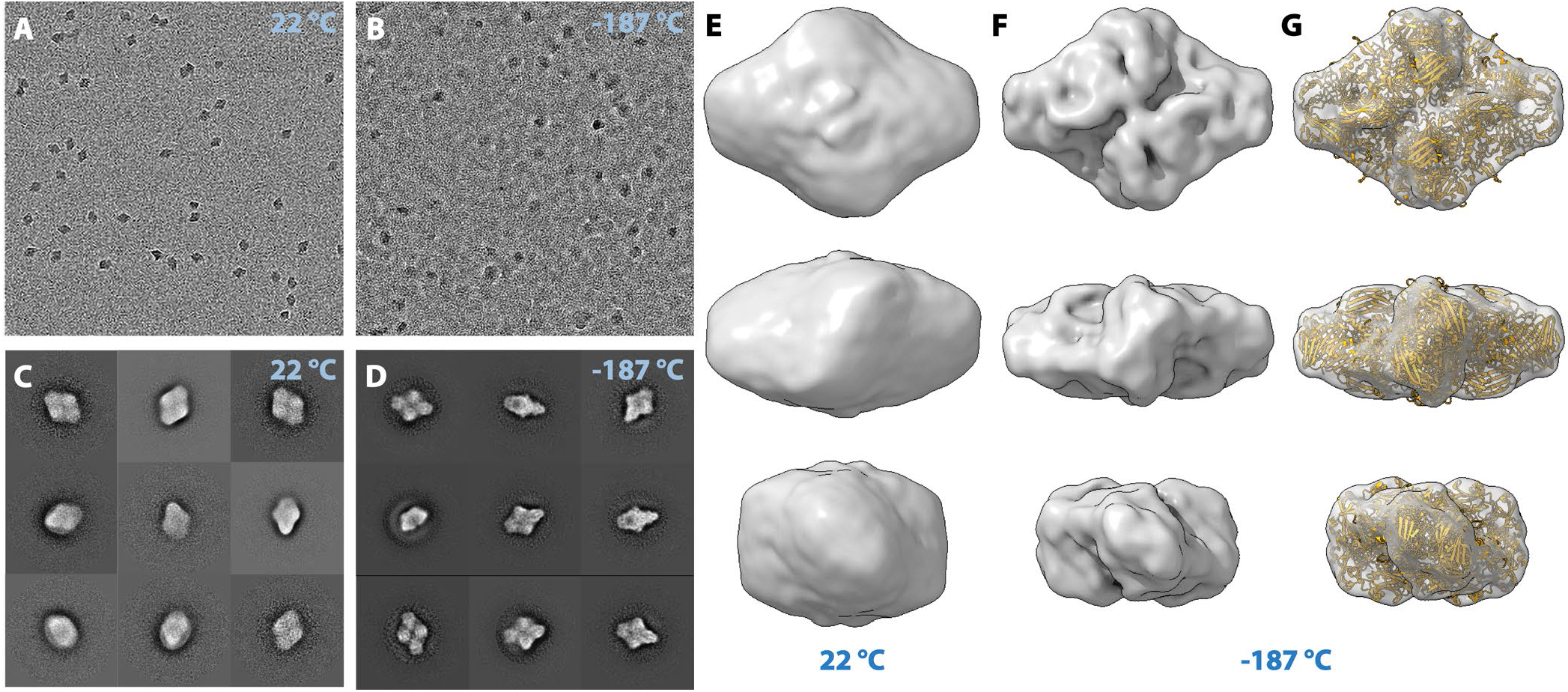
Cryo-electron microscope images and reconstructions of room temperature versus cryogenically landed β-galactosidase cations. Panels A and B present representative micrographs of either room temperature or cryo-landed β-Galactosidase cations. Panels C and D present the 2D class averages that resulted from either the room temperature or cryo-landing specimens, respectively. Panel E shows the 3D reconstruction that resulted from the room temperature landing, whilst panel F presents the structure obtained from the cryo-landing experiment. Panel G shows the fit of the cryo-landed model with the known high resolution ribbon structure.

**Figure 3.**
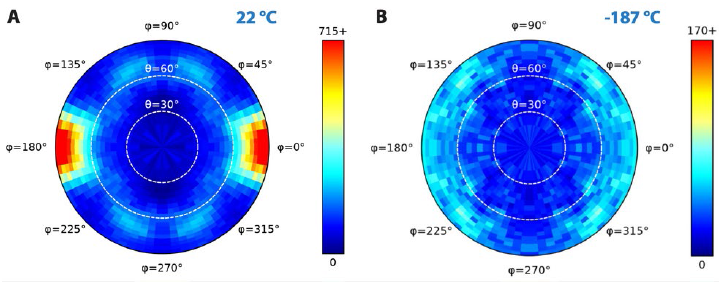
Comparison of angular particle orientations resulting from either room temperature (A) or cryogenically (B) landed β-Galactosidase cations. Note the room temperature landing sample resulted in a highly preferred orientation while the cryo-landed sample does not.

Panels A and B of **Figure 4** display representative micrographs of the resulting room temperature and cryo-landed proteasome particles. As in the previous β-Galactosidase experiment, the quality of the particles cryogenically landed on ice is substantially better than those landed at room temperature, with more detail apparent in both the 2D class averages (panels C and D) and the 3D reconstruction (panels E and F). Also mirroring the previous experiment, the results suggest protection from degradation could be improved: despite demonstrating some protection with the cold, amorphous ice-coated surface, the quality of the cryogenically landed particles is still lower than would be expected in a conventionally prepared proteasome specimen. Interestingly, several of the 2D class averages of the cryo-landed proteasome cations show what appear to be droplets or water build-up around the edges. Note mass spectral measurements were acquired (data not shown) and do not indicate the presence of sufficient water molecules to explain the droplets. We conclude that the water must build up around the particles post-deposition. Indeed, this effect points us back to the β-Galactosidase experiment, where it was much less pronounced but still present as demonstrated by the middle-left class average of **Figure 2D**. At present the mechanism of this drop formation is unclear and is the continued subject of investigation. Unlike the β-Galactosidase experiment, there is no obvious difference when comparing the angular orientation distributions of the proteasome particles in both the room temperature and cryogenically landed experiments. Both samples exhibit the classic preferential orientation towards top and side views seen in traditional proteosome reconstructions. It is unclear whether this is a sample-specific effect or related to the different nature of ice formation in the two experiments.

**Figure 4.**
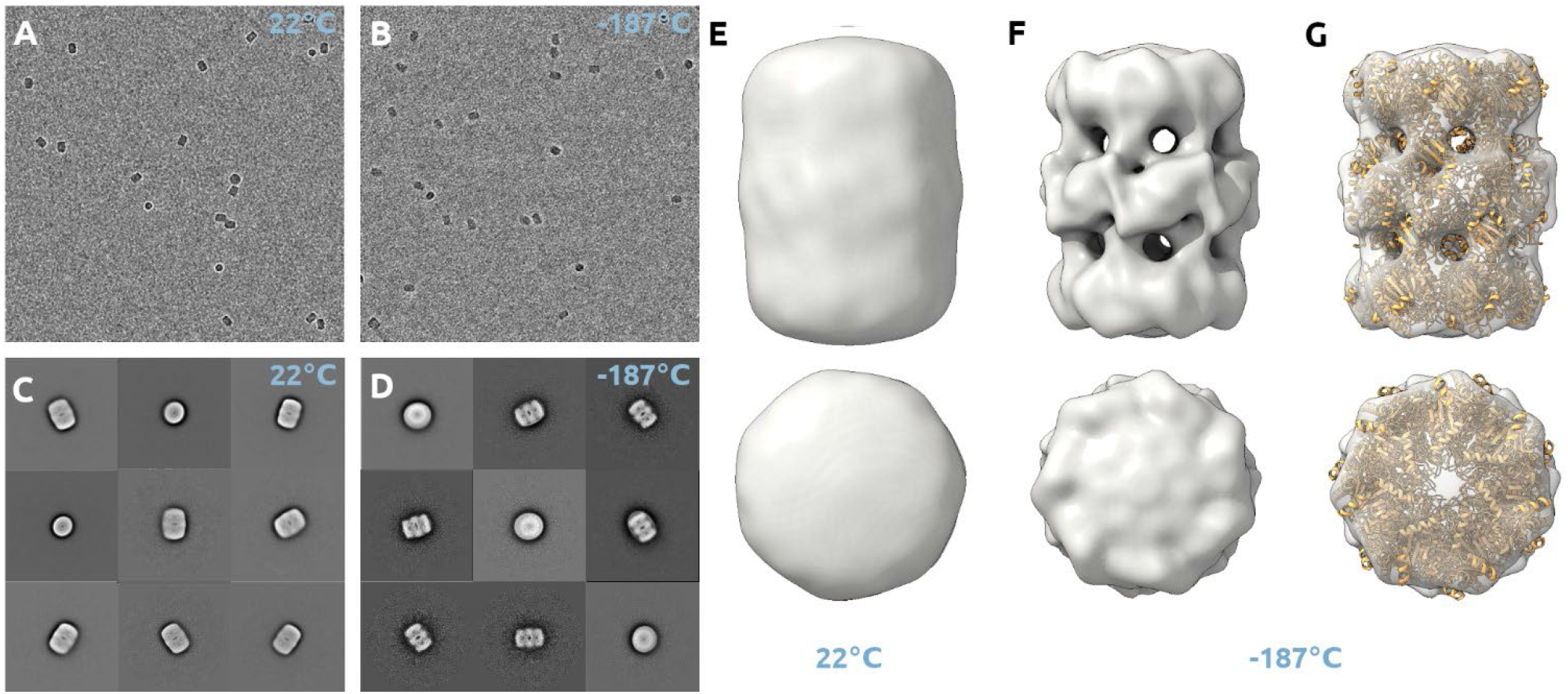
Cryo-electron microscope images and reconstructions of room temperature versus cryogenically landed proteasome cations. Panels A and B present representative micrographs of either room temperature or cryo-landed proteasome cations. Panels C and D present the 2D class averages resulting from these respective landings. Panel E shows the 3D reconstruction that resulted from the room temperature landing, whilst panel F presents the structure obtained from the cryo-landing experiment. Panel G shows the fit of the cryo-landed model with the known high resolution ribbon structure.

## DISCUSSION

Recently, we and others have reported on the modification of a hybrid Orbitrap mass spectrometer to permit the landing of protein– protein complex cations onto TEM grids at room temperature. In our initial experiments, the deposition of particles onto bare carbon TEM grids with negative stain TEM analysis showed that the particles had degraded. Recent work by Rauschenbach et al. confirmed this degradation by imaging room temperature-landed particles, subsequently plunged into liquid nitrogen and kept cold for imaging.^14^ The resulting 3D reconstructions of β-Galactosidase appear comparable to those shown in **Figure 2** here. Our previous work addressed the problem of particle degradation by use of a chemical landing matrix, which allowed for the highest resolution attainable using negative stains.^10, 11^ Unfortunately, this matrix is incompatible with cryo-EM, which would afford a higher resolution. In this work, we describe a novel cryogenic landing apparatus that allows us to explore the effects of a cryogenic landing surface for the preservation of particles and the formation of thin films of amorphous ice in vacuo. With this device, we demonstrate considerable advantages with respect to particle preservation over room temperature deposition. And because the cryogenic landing of the particles presumably captures them in their native gas-phase conformation, this work represents the most direct measurement of the gas-phase structure of protein–protein complex cations as they traverse the mass spectrometer collected to date.

There are several possibilities for why cryo-landing better preserves particle structure and can improve the distribution of particle orientations. First, the cryogenic temperature may prevent further dehydration of the protein–protein complex cation whilst also slowing the rate of deleterious surface interactions. Second, the thin film of ice could be partially serving the protective role of the chemical matrix we recently described. Third, although the role ice plays in radiation damage is not currently well understood, it is possible that embedding the particles in the ice present in cryo-landed samples leads to less radiation damage by the electron beam.^20^ Finally, in the case of β-Galactosidase we observe more orientations in the cryo-landed data shown here, presumably due to the reduced particle– carbon surface interactions.

Despite these considerable improvements over room temperature landing, we have not yet achieved structural resolution comparable to that of conventional plunge-frozen cryo-EM samples. We believe there are five potential drivers causing apparent structural degradation that future work will seek to address. First, as particles transition through the mass spectrometer, it is possible that dehydration is degrading their structural integrity, precluding higher resolution reconstructions even if the particles are perfectly preserved when landed on the grid surface. Second, cooling the surface may not be sufficient to completely prevent the vacuum exposure and/or the surface itself from inducing structural degradation. Third, structural damage may be induced upon deposition due to suboptimal deposition energy. Fourth, the presence of the droplets, observed most prominently with the proteasome sample, could be problematic in ways we cannot yet understand. Finally, it is possible that interactions of the charged particles with the surface could be causing structural degradation.^21-23^

Ongoing work in our group aims to address these potential challenges. To limit dehydration of particles, we are currently examining modifications to how we generate the ions and are working towards freezing the particles whilst still in the gas phase.^24^ The goal of gasphase freezing is to create amorphous ice-coated particles directly after entry into the mass spectrometer—particles that can land on the cryogenically cooled grid already protected from vacuum and degradation. Our second approach will be to continue exploring chemical matrices and grid surface modifications so that our matrix-landing concept can be compatible with the requirements of cryo-EM. Moving forward, if we can control the amount of matrix to allow for the particles’ protection as well as cryo-EM imaging, we can combine the benefits of both techniques. Finally, there are many parameters of water and particle co-deposition yet to be explored—including timing (water first, during, after, etc.), use of a water molecular beam versus water vapor, the rate of water deposition, the temperature of deposition, among many others.

In summary, we describe here a cryogenic landing apparatus that allows for temperature control of a TEM grid within a mass spectrometer environment, formation of amorphous ice, and a means to remove the sample from vacuum without contamination. With this device, we demonstrate considerably improved structural resolution, as evidenced by cryo-EM imaging of cryogenically landed protein–protein complex cations. In addition to improving structural quality, this process also provided a much more diverse particle orientation than room temperature landing. We believe the future work outlined above will provide a path for continued improvements that ultimately allow for the direct coupling of mass spectrometry with cryo-EM.

## Supporting information

FSC curves.

## ASSOCIATED CONTENT

### Supporting Information

The Supporting Information is available free of charge on the ACS Publications website.

FSC curves for β-Galactosidase and the 20S Proteasome.

## AUTHOR INFORMATION

### Authors

Michael S. Westphall-Department of Biomolecular Chemistry, University of Wisconsin-Madison, Madison, Wisconsin 53715, United States;

Kenneth W. Lee-Department of Biomolecular Chemistry, University of Wisconsin-Madison, Madison, Wisconsin 53715, United States;

Colin A. Hemme-Department of Chemistry, University of Wisconsin-Madison, Madison, Wisconsin 53715, United States;

Austin Z. Salome-Department of Chemistry, University of Wisconsin-Madison, Madison, Wisconsin 53715, United States;

Keaton Mertz-Department of Chemistry, University of Wisconsin-Madison, Madison, Wisconsin 53715, United States;

## Author Contributions

MSW and JJC conceptually designed the research; MSW and KM constructed the instrumentation; MSW, KWL, AZS, CH and TG performed the experiments; TG conducted 3D structural reconstructions; MSW, TG, and JJC wrote the paper.

## ACKNOWLEDGMENT

We are grateful for support of this project by the National Institutes of Health R35GM118110 grant (to JJC), the Morgridge Institute for Research, and the University of Wisconsin-Madison. We are grateful to Yifan Cheng and Zanlin Yu for generously providing the 20S proteasome core particles. We thank Alicia Williams for her literary support in drafting this manuscript.

