## Supplementary material for "Cryogenic Soft Landing Improves Structural Preservation of Protein Complexes": FSC curves.

### Matrix-Landing Mass Spectrometry for Electron Microscopy Imaging of Native Protein Complexes

### Table of Contents

SUPPLEMENTARY FIGURE 1 .......................................................................2

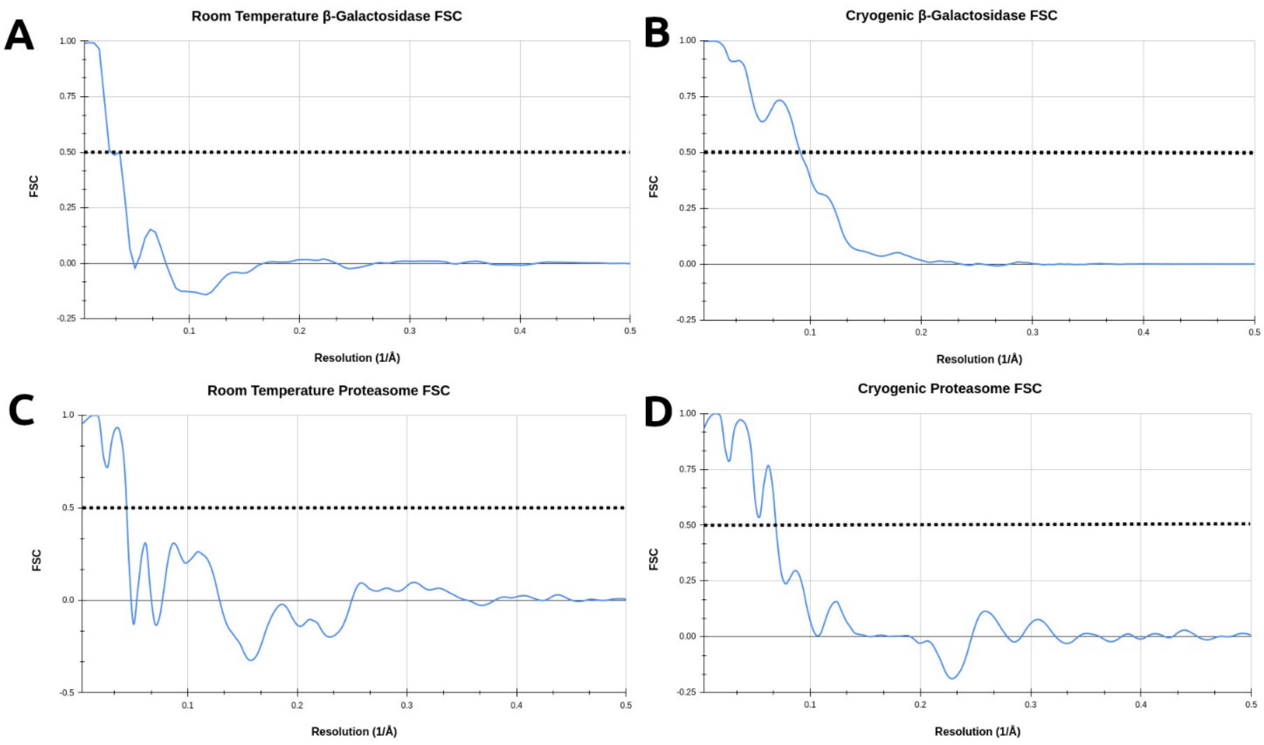


**Supplementary Figure 1.** FSC curves for β-Galactosidase and the 20S Proteasome core particle. Curves were generated using Compute FSC, a program within cisTEM that calculates a FSC between two maps. The β-Galactosidase maps were correlated to a map of the PDB 1F4H simulated at 2 Å. The 20S Proteasome maps were correlated to a map of the PDB 3J9I also simulated at 2 Å resolution. These calculations also included a mask generated from the PDB's simulated at 15 Å and then sharpened with a positive B-factor to remove hard edges. (A) Room Temperature β-Galactosidase FSC curve with a resolution of ~80 Å. (B) Cryogenic β-Galactosidase FSC curve with a resolution of ~28 Å. (C) Room Temperature 20S Proteasome FSC curve with a resolution of ~55 Å. (D) Cryogenic 20S Proteasome FSC curve with a resolution of ~37 Å.
